# Synergistic Arg-C Ultra and Lys-C Digestion for Quantitative Proteomics

**DOI:** 10.1101/2025.07.15.664461

**Authors:** Vyas Pujari, Joseph Crapse, Connor Nisbet, Gloria Bao, Wessley Ferguson, Christopher M Hosfield, Michael Rosenblatt, Felix Keber, Martin Wühr

## Abstract

Shotgun proteomics hinges on complete enzymatic digestion of proteins into peptides. Incomplete digestion narrows proteome coverage and inflates variability in quantitative workflows, whether label-free DIA or multiplexing with isobaric tags. Sequential Lys-C/Trypsin digestions mitigate missed cleavages at lysine residues, but arginine sites remain a persistent challenge. Arg-C Ultra, a recently released cysteine protease, efficiently targets arginine residues but requires reducing conditions that inactivate Lys-C activity and compromise NHS-ester labeling in multiplexed workflows. Here, we systematically characterized Arg-C Ultra and Lys-C with chromogenic substrates that mimic arginine- and lysine-containing peptides, as well as shotgun proteomics. Arg-C Ultra operates optimally at room temperature, pH 7.5–8.5, under reducing conditions, whereas Lys-C is most active at 37 °C, pH 7.5–8.5, yet rapidly loses activity when exposed to common reductants. Among tested reducing agents, 1 mM TCEP uniquely preserved TMTpro integrity while sustaining Arg-C Ultra activity. Guided by these insights, we established a sequential digestion workflow that is fully compatible with both label-free DIA and TMTpro multiplexing. Proteins are first digested overnight with Lys-C at 37 °C (pH 8.5), then treated with 1 mM TCEP and Arg-C Ultra at room temperature (pH 8.5). The resulting peptides can be analyzed directly by label-free DIA or subjected to TMTpro labeling for multiplexed quantification. Applied to HeLa cell lysates, this protocol achieved >99% arginine and 95% lysine cleavage efficiencies, boosting the number of quantified proteins by 6% in label-free DIA and 11% in TMTproC experiments. Replicate measurements displayed reproducibility that approached the limits set by ion statistics. Thus, the introduced synergistic Lys-C/Arg-C Ultra digestion strategy enhances proteome coverage with excellent quantitative reproducibility across both label-free and multiplexed platforms.

## Introduction

Shotgun proteomics has revolutionized our ability to comprehensively analyze cellular proteomes, enabling quantitative comparisons across various biological conditions^1,2^. The success of these approaches fundamentally depends on efficient enzymatic digestion to convert complex protein mixtures into peptides suitable for liquid chromatography-mass spectrometry analysis^3^. Incomplete or variable digestion introduces biases that limit both proteome coverage and quantitative accuracy and precision^4–6^, making digestion optimization a promising avenue for improving proteomics workflows.

Trypsin is the most widely used protease in proteomics, cleaving after arginine and lysine residues to generate peptides that typically carry a charge on the N-terminal alpha-amino group and on the arginine or lysine at the C-terminus^7,8^. The resulting doubly charged (2+) peptides are generally optimal for shotgun proteomics analysis^9,10^. However, while trypsin has great selectivity, it often misses cleavage sites^11,12^. As a result, state-of-the-art protein digestion protocols commonly use a combination of trypsin and Lys-C^13,14^. Lys-C helps address trypsin’s high missed-cleavage rate after lysine residues, but relatively high missed-cleavage rates after arginine remain^12,15^.

Figure 1 summarizes a commonly used bottom-up proteomics workflow using Lys-C/Trypsin digestion including SP3 magnetic bead precipitation for protein cleanup^14,16,17^. Following cell lysis and protein precipitation, each condition of interest is prepared separately with bead-bound proteins initially digested with the highly denaturant-tolerant^18^ Lys-C in 2 M guanidine hydrochloride, then subjected to a second digestion with both Lys-C and trypsin after dilution to a milder 0.5 M guanidine hydrochloride (Figure 1a). For label-free analysis, samples from each condition are purified independently via C18 cleanup. For multiplexed proteomics, the resulting peptides from each condition are labeled with a unique TMTpro isobaric tag, combined, and purified via C18 cleanup.

**Figure 1.**
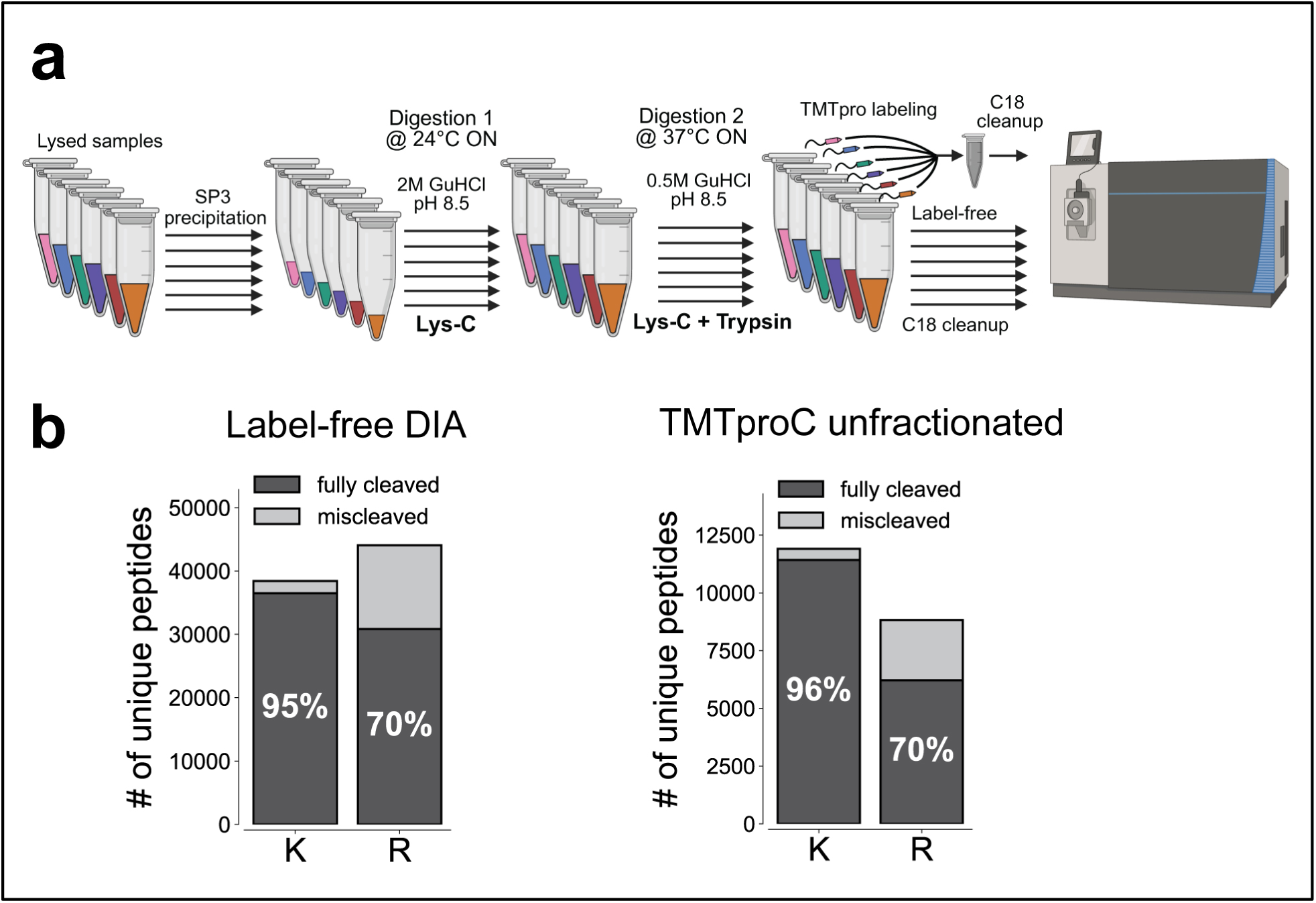
Standard Lys-C/Trypsin workflow and associated cleavage efficiency in label-free and multiplexed proteomics workflows. **(a)** Schematic overview of a standard Lys-C/Trypsin workflow. Lysates undergo protein cleanup using SP3 precipitation. Bead-bound proteins are initially digested overnight with Lys-C (20 ng/uL) in 2 M GuHCl, 10 mM EPPS (pH 8.5) at 24 °C. The following day, the GuHCl concentration is diluted to 0.5 M, and digestion proceeds overnight at 37 °C with an additional Lys-C (20 ng/uL) and Trypsin (10 ng/uL). For label-free experiments, samples are separately desalted via C18 cleanup and analyzed on the mass spectrometer. For multiplexed analysis samples are vacuum-evaporated, resuspended in 200 mM EPPS (pH 8.0), labeled with TMTpro, combined, and desalted. Created with BioRender.com. **(b)** Proteomics analysis of HeLa lysate via TMTproC and label-free DIA reveals cleavage efficiencies from the standard workflow, showing ∼5% of peptides where the first basic amino acid is lysine (K) containing a missed cleavage, and 30% of peptides where the first basic amino acid is arginine (R) containing a missed cleavage.

Despite widespread adoption, standard Lys-C/Trypsin protocols exhibit persistent limitations in cleavage efficiency. Missed cleavage rates after lysine remain relatively low (∼5%), whereas peptides containing arginine show substantially higher rates (∼30%) in both label-free DIA and multiplexed analyses (Figure 1b). This disparity represents a potential source of analytical variability and limits the depth of proteome coverage achievable in quantitative proteomic experiments. Improved cleavage efficiency can help collapse peptide variants, leading to stronger signals and better distinction between peptides and background noise. Additionally, peptides containing missed cleavages often have higher charge states, which makes peptide identification and quantification more challenging^10,19–21^.

Promega recently released Arg-C Ultra, a cysteine protease isolated from *Porphyromonas gingivalis* (also known as gingipain R2 or RgpB) with high activity and specificity for arginine residues. Originally characterized as an arginine-specific virulence factor from this periodontal pathogen^22–26^, this enzyme presents an opportunity to address the main source of missed cleavages in standard proteomics sample preparation workflows. At first glance, Arg-C Ultra exhibits higher catalytic activity and broader pH tolerance than the well-established—yet seldom used—standard Arg-C (clostripain)^27,28^. However, Arg-C Ultra requires reducing conditions to retain activity, which creates potential incompatibilities with established proteomics workflows. For example, NHS-ester based isobaric labeling reagents such as TMTpro are susceptible to hydrolysis by reducing agents. We also found that reducing conditions compromise the stability of Lys-C. Additionally, the commonly used denaturant guanidine inhibits Arg-C Ultra, likely due to its structural similarity with arginine side chains resulting in competitive inhibition. While the combination of Arg-C Ultra and Lys-C offers, in principle, an attractive path to highly efficient trypsin-like digestion, successful implementation must address these constraints.

Here, we systematically characterize the biochemical properties and optimal reaction conditions for both Arg-C Ultra and Lys-C using chromogenic substrate assays and validate our findings via shotgun proteomics. We demonstrate that careful selection of reducing agents and reaction conditions enables the development of a sequential Lys-C/Arg-C Ultra digestion protocol that dramatically improves arginine cleavage efficiency while maintaining compatibility with both label-free and multiplexed proteomics workflows. Our resulting protocol achieves >99% arginine cleavage efficiency and provides substantial improvements in proteome coverage for both label-free DIA and TMTpro-multiplexed proteomics experiments.

## Results and Discussion

### Characterization of Arg-C Ultra, a novel cysteine protease

To determine optimal conditions for Arg-C Ultra activity, we characterized the enzyme’s biochemical properties using both UV–Vis-based chromogenic substrate assays and shotgun proteomics. Specifically, we examined how denaturants, pH, temperature, and reducing agents affect its performance. In the UV-Vis assays, the chromogenic substrate Nα-Benzoyl-L-arginine 4-nitroanilide was used, which produces a measurable increase in absorbance at 385 nm upon cleavage (Figure 2a, b). We first used this assay to evaluate Arg-C Ultra activity in the presence of various denaturants. Denaturants are often used during digestion to unfold proteins and enable their digestion^29,30^. However, higher denaturant concentration often also inhibits protease activity^31,32^. We observed drastic inhibitory effects of guanidine hydrochloride on Arg-C Ultra activity even at relatively low concentrations (50 mM) (Figure 2c), which is likely explained by the structural similarity of guanidine and arginine. When we tested Arg-C Ultra activity in presence of urea, we observed decreased enzyme activity with increased urea concentration in chromogenic substrate assays (Figure 2d). Similarly, the number of unique, fully cleaved peptides detectable in shotgun proteomics experiments decreased with higher urea concentrations (Figure 2e). However, while higher urea concentrations lead to decreased enzyme activity, it is possible that a denaturant is needed to unfold some substrates and make them digestible; the assays used so far might not be able to detect this. To address this question, we analyzed *E. coli* lysate before and after digestions with Coomassie-stained gels. Surprisingly, protein digestion was most complete in the absence of urea; however, undigested protein bands were present at all tested urea concentrations from 0 to 6 M (Figure 2f, example indicated by red arrow). These proteins may lack arginine residues or possess arginines that remain inaccessible to enzymatic digestion, even under the denaturing conditions tested here. Besides having no apparent advantage in the assays used here, urea is additionally unfavorable due to its decomposition products: isocyanate, which reacts with peptide N-termini and lysine residues to form carbamylated adducts, and ammonia, which can act as a nucleophile and react with TMTpro NHS-esters^33^. Carbamylation impairs peptide identification and also blocks residues required for isobaric labeling^34^. Thus, we find that not using any denaturants appears optimal for both Arg-C Ultra activity and compatibility with TMTpro labeling. Next, we evaluated Arg-C Ultra’s activity dependence as a function of pH. We find that Arg-C Ultra maintains robust activity across a broad pH range, with peak performance between pH 7.5–8.5 in both chromogenic assays and shotgun proteomics experiments (Figure 2g-h). Lastly, we found that Arg-C Ultra incubation at room temperature outperformed digestion at 37 °C (Figure 2i), which may be due to higher enzyme stability at lower temperatures. Notably, under these room temperature conditions with 1 mM TCEP, less than 1% of peptides contained a missed cleavage. These findings establish optimal conditions for Arg-C Ultra as pH 7.5–8.5 at room temperature in the absence of denaturants, with reducing conditions required for activity.

**Figure 2.**
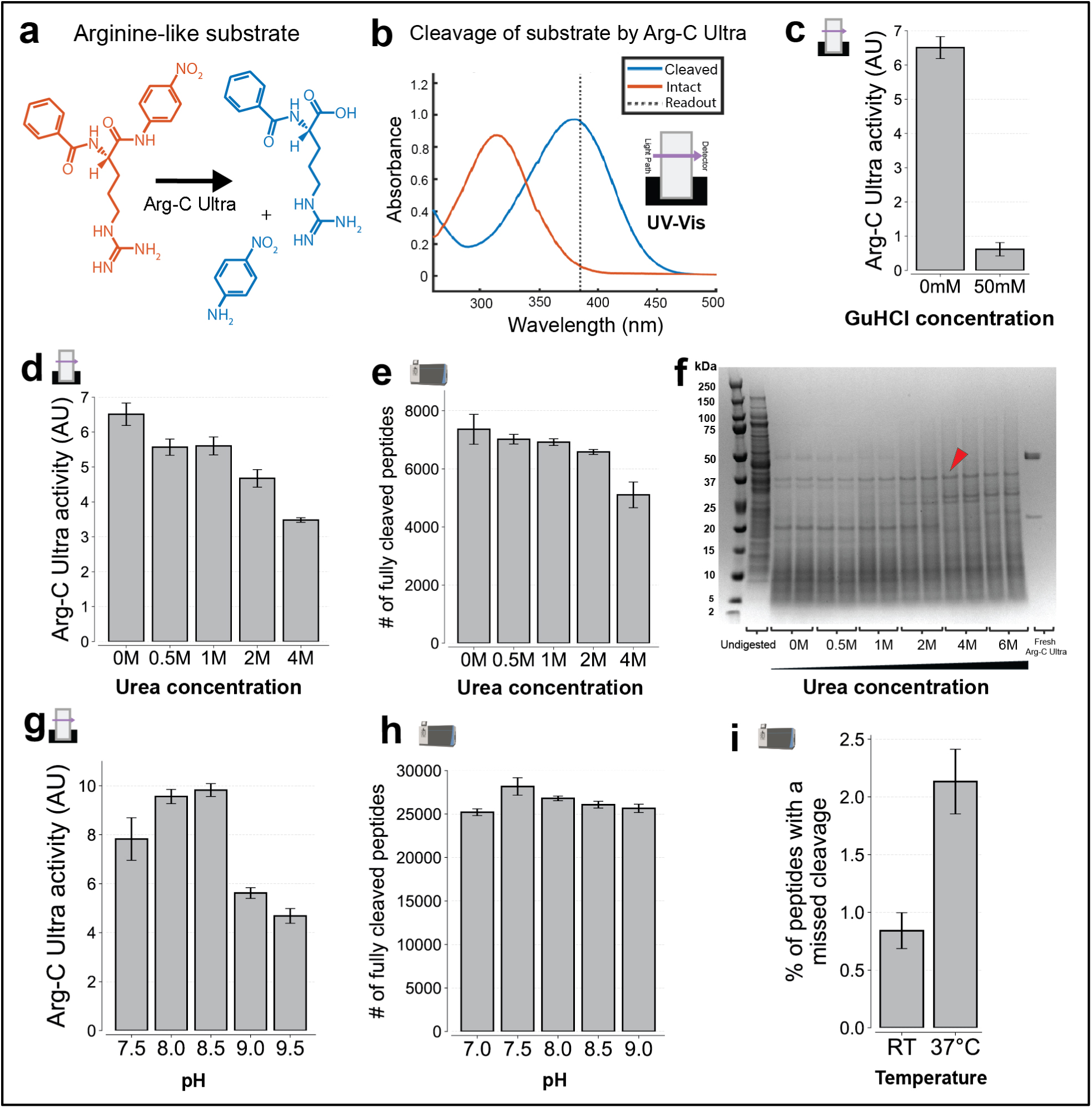
Characterization of Arg-C Ultra protease activity and conditions affecting its performance. Icons indicate experiment type: shotgun proteomics or UV-Vis. **(a)** Structure of arginine-like chromogenic substrate before and after cleavage by Arg-C Ultra, used in UV- Vis activity assays. **(b)** UV-Vis absorbance spectra of intact and cleaved substrate. Cleaved substrate shows strong absorbance at 385 nm, while the intact substrate does not. Enzyme activity was measured by the increase in absorbance at 385 nm. **(c)** Effect of guanidine on Arg-C Ultra activity, assessed via UV-Vis. Even a relatively low GuHCl concentration (50 mM) significantly reduces Arg-C Ultra activity, likely due to structural similarity between guanidine and arginine. **(d)** Effect of urea concentration on Arg-C Ultra activity, assayed via UV-Vis. Activity is highest with no urea and diminishes with increasing urea concentration. **(e)** Effect of urea concentration on digestion by Arg-C Ultra, assessed by shotgun proteomics. *E. coli* lysate was digested by Arg-C Ultra at pH 8.5 overnight at room temperature, using varying urea concentrations. The number of unique, fully cleaved peptides was highest with no urea and diminishes with increasing urea concentration. **(f)** Coomassie-stained gels showing residual proteins after digestion by Arg-C Ultra across varying urea concentrations. Samples were prepared as in (e), except proteins still bound to the beads were additionally released. Undigested proteins persist at all conditions (example shown with red arrow), though the fewest high-molecular-weight bands appear with no urea. **(g)** Effect of pH on Arg-C Ultra activity, assayed via UV-Vis. Arg-C Ultra is active across the tested pH range, peaking near pH 8.0–8.5 **(h)** Effect of pH on digestion by Arg-C Ultra, assessed by shotgun proteomics. HeLa lysate was digested with Arg-C Ultra at 37 °C in the absence of urea, using 20 mM EPPS buffer containing 1 mM TCEP at various pH values. pH 7.5–8 yielded the highest number of unique, fully cleaved peptides. **(i)** Effect of temperature on digestion by Arg-C Ultra, assessed by shotgun proteomics. Same as (h), but digested overnight at pH 8.5 either at room temperature or at 37 °C. Cleavage efficiency was higher at RT. Error bars represent ±1 standard deviation; n = 3 for all UV-Vis experiments and n = 2 (independent digestions) for all proteomics experiments. All shotgun proteomics data were acquired using label-free data-dependent acquisition (DDA).

### Low TCEP concentration is compatible with TMTpro labeling

The requirement for reducing conditions for Arg-C Ultra activity also raised concerns about compatibility with NHS-ester based isobaric labeling, as common reducing agents like β- mercaptoethanol (BME) and dithiothreitol (DTT) can hydrolyze NHS-ester bonds and impair labeling efficiency (Figure 3a). For compatibility with multiplexed proteomic workflows, this challenge must be resolved. We systematically evaluated the effect of various reducing agents on TMTpro NHS-ester stability using UV-Vis spectroscopy, monitoring the characteristic absorbance change at 260 nm that accompanies NHS-ester hydrolysis (Figure 3a-b). Comparative analysis revealed dramatic differences between reducing agents. Both BME and DTT at 1 mM concentrations caused rapid and extensive NHS-ester hydrolysis within minutes (Figure 3c). In contrast, TCEP at the same concentration caused substantially less hydrolysis, though the effect was concentration-dependent, with higher TCEP concentrations (>5 mM) showing increased interference (Figure 3d). To validate these findings in a practical context, we performed TMTpro0 labeling experiments with HeLa peptides in the presence of varying TCEP concentrations. Proteomics analysis confirmed that 1 mM TCEP had minimal impact on labeling efficiency, achieving comparable performance to TCEP-free conditions, while 5 mM TCEP did slightly reduce labeling efficiency (Figure 3e). These results established 1 mM TCEP as a robust choice to preserve TMTpro labeling compatibility while maintaining Arg-C Ultra activity.

**Figure 3.**
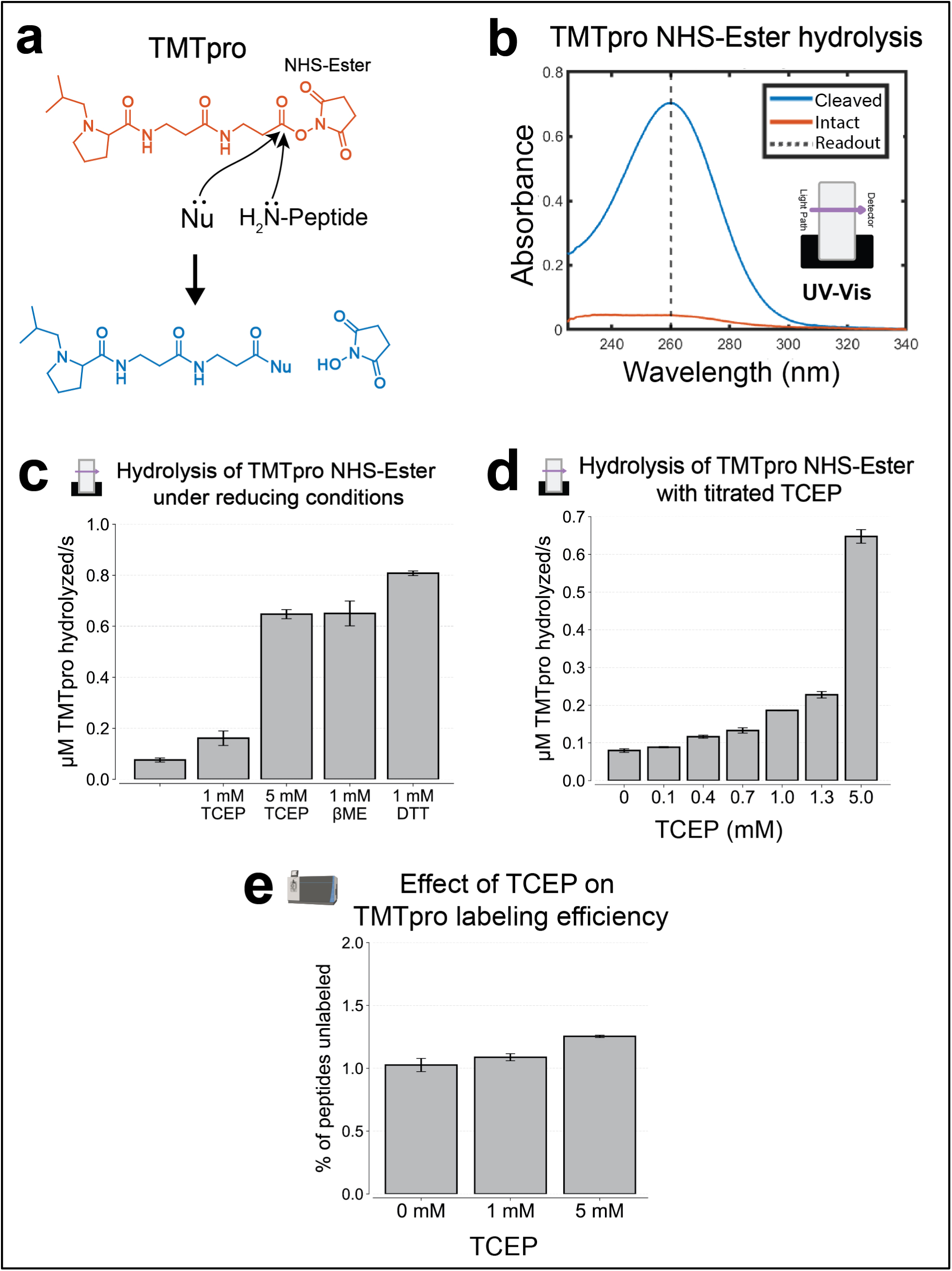
Compatibility of low TCEP concentrations with NHS-Ester-based isobaric labeling. Icons indicate experiment type: shotgun proteomics or UV-Vis. **(a)** Overview of nucleophilic (Nu) hydrolysis susceptibility of isobaric tags like TMTpro NHS-Ester. Common reducing agents (e.g., β-mercaptoethanol (BME), dithiothreitol (DTT)) can hydrolyze NHS- esters, potentially impairing TMTpro labeling efficiency. **(b)** UV-Vis absorbance spectra of intact and hydrolyzed TMTpro0 NHS-ester. Intact TMTpro0 weakly absorbs at 260 nm, while hydrolyzed strongly absorbs. Absorbance at 260 nm was converted to concentration using Beer’s Law (ε = 9700 M⁻¹cm⁻¹)^36^. **(c)** Comparative assessment of NHS-ester hydrolysis by reducing agents BME, DTT, and TCEP. Without reducing agents, spontaneous hydrolysis occurs due to hydroxide ion interactions. DTT and BME (1 mM) rapidly induce extensive NHS-ester hydrolysis, whereas TCEP at the same concentration causes substantially less hydrolysis. **(d)** UV-Vis assay showing increased TMTpro NHS-ester hydrolysis with rising TCEP concentrations. **(e)** Effect of TCEP on TMTpro Labeling Efficiency. Shotgun proteomics analysis showing that HeLa peptides labeled with TMTpro0 in the presence of 1 mM TCEP achieve labeling efficiencies comparable to TCEP-free conditions, while 5 mM TCEP reduces labeling efficiency, increasing the fraction of unlabeled peptides by 23%. TMTpro0 was searched as a variable modification on lysine residues and peptide N-termini. Error bars represent ±1 standard deviation; n = 3 for all experiments. For proteomics experiments, each replicate was labeled independently. Shotgun proteomics data were acquired using a TMTpro-SPS-MS3 method.

### Characterization of Lys-C reveals incompatibility with reducing conditions required for Arg-C Ultra

We characterized *Lysobacter*-derived Lys-C using a similar approach to Arg-C Ultra, employing both chromogenic substrate assays and shotgun proteomics, which revealed distinctly different optimal conditions compared to Arg-C Ultra. In the UV-Vis activity assays, we used the chromogenic substrate Nα-Benzyloxycarbonyl-L-lysine thiobenzyl ester and measured enzyme activity by monitoring the reduction in absorbance at 240 nm upon cleavage (Figure 4a-b). Unlike Arg-C Ultra, Lys-C showed tolerance to urea in UV-Vis activity assays, maintaining comparable activity across 0-4M urea concentrations (Figure 4c). However, shotgun proteomics experiments revealed that the number of unique, fully cleaved peptides was still highest in the absence of urea, in accordance with existing findings^35^ (Figure 4d). Protein digestion patterns analyzed by Coomassie staining showed no clear advantage at higher urea concentrations: some bands disappeared while others appeared, with undigested proteins present at all tested concentrations from 0 to 6 M (Figure 4e, example indicated by red arrow). pH optimization showed that Lys-C activity was highest at pH 8.0–8.5 in chromogenic substrate assays and that pH 7.0–8.0 maximized the number of unique, fully cleaved peptides in shotgun proteomic experiments (Figure 4f-g). We suspect that the higher numbers of peptides identified with shotgun proteomics at lower pH might stem from the fewer background side-reactions, like oxidation, that might make it easier to identify canonical peptides. Temperature studies using proteomics analysis confirmed that 37 °C was optimal for digestion efficiency (Figure 4h), possibly due to favorable enzyme kinetics at elevated temperature.

**Figure 4.**
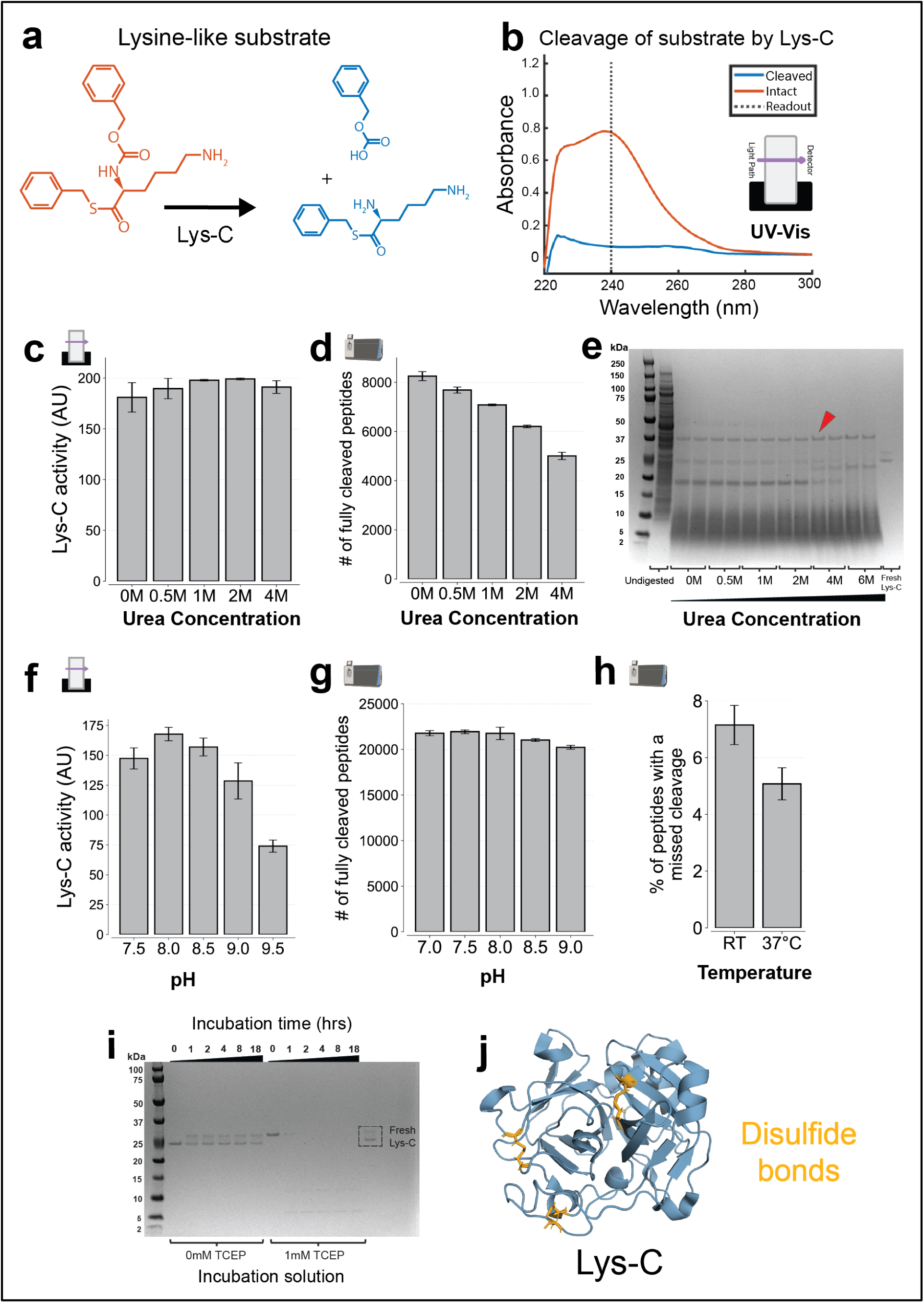
Characterization of Lys-C protease activity and conditions affecting its performance. Icons indicate experiment type: shotgun proteomics or UV-Vis. **(a)** Structure of lysine-like chromogenic substrate before and after cleavage by Lys-C, used in UV-Vis activity assays. **(b)** UV-Vis absorbance spectra of intact and cleaved substrate. Intact substrate shows strong absorbance at 240 nm, which decreases upon cleavage. Enzyme activity was measured by the reduction in absorbance at 240 nm. **(c)** Effect of urea concentration on Lys-C activity, assayed via UV-Vis. Activity is invariant of urea concentration up to 4M. **d)** Effect of urea concentration on digestion by Lys-C, assessed by shotgun proteomics. *E. coli* lysate was digested by Lys-C at pH 8.5 overnight at 37 °C, using varying urea concentrations. The number of unique, fully cleaved peptides was highest with no urea and diminishes with increasing urea concentration. **e)** Coomassie-stained gels showing residual proteins after digestion by Lys-C across varying urea concentrations. Samples were prepared as in (d), except proteins still bound to the beads were additionally released. Undigested proteins persist at all conditions (example shown with red arrow). **f)** Effect of pH on Lys-C activity, assayed via UV-Vis. Activity peaks near pH 8.0. **g)** Effect of pH on digestion by Lys-C, assessed by shotgun proteomics. HeLa lysate was digested with Lys-C at 37 °C in the absence of urea, using 20 mM EPPS buffer at various pH values. pH 7–8 yielded the highest number of unique, fully cleaved peptides. **h)** Effect of temperature on digestion by Lys-C, assessed by shotgun proteomics. Same as (g), but digested overnight at pH 8.5 either at room temperature or at 37 °C. Cleavage efficiency was higher at 37°C. **(i)** TCEP results in degradation of Lys-C. Lys-C was incubated with or without 1 mM TCEP in 20 mM EPPS (pH 8.5) and analyzed by Coomassie-stained gel at various time points. Without TCEP, Lys-C is stable, displaying two bands—likely the oxidized and reduced variants. In the presence of 1 mM TCEP, only the reduced band is visible, which appears to degrade in as little as 1 hour. **(j)** Crystal structure of Lys-C indicates disulfide bonds (PDB ID: 4NSY). Disulfide bonds can be disrupted under reducing conditions. Error bars represent ±1 standard deviation; n = 3 for all UV-Vis experiments and n = 2 (independent digestions) for all proteomics experiments. All shotgun proteomics data were acquired using label-free data-dependent acquisition (DDA).

Given that Arg-C Ultra is a cysteine protease requiring reducing agents, we tested whether such reducing agents might affect Lys-C activity. Surprisingly, we found that Lys-C activity was rapidly inhibited in the presence of 1mM TCEP (Figure S1a). Furthermore, Lys-C pre-incubated in TCEP lost its ability to digest cell lysate (Figure S1b). Analyzing Lys-C before and after exposure to TCEP on a Coomassie-stained gel showed an upward shift followed by rapid degradation, with significant protein breakdown observable within one hour (Figure 4i). Examination of the Lys-C crystal structure revealed multiple disulfide bonds (Figure 4j). We believe that upon TCEP exposure, these disulfide bond(s) are broken, leading to an upward shift in the Coomassie gel. We suspect that the broken disulfide bonds make the protein less stable and prone to proteolysis, thereby inactivating Lys-C under reducing conditions. Unfortunately, this finding points towards a fundamental incompatibility between activity requirements for Lys-C and Arg-C-Ultra. To overcome this limitation, we opted to employ sequential digests each geared towards the respective enzymes.

### Sequential Lys-C/Arg-C Ultra digestion significantly improves proteome coverage for TMTpro multiplexed and label-free DIA workflows

Based on the individual enzyme characterizations, we developed a sequential digestion protocol that leverages the optimal conditions for each enzyme while addressing their incompatibilities. Proteins are first digested with Lys-C at 37 °C under non-reducing conditions (20 mM EPPS) overnight, followed by addition of 1 mM TCEP and Arg-C Ultra for a second overnight incubation at room temperature (Figure 5a). This approach allows Lys-C to function under optimal conditions before introducing the reducing environment required for Arg-C Ultra.

**Figure 5:**
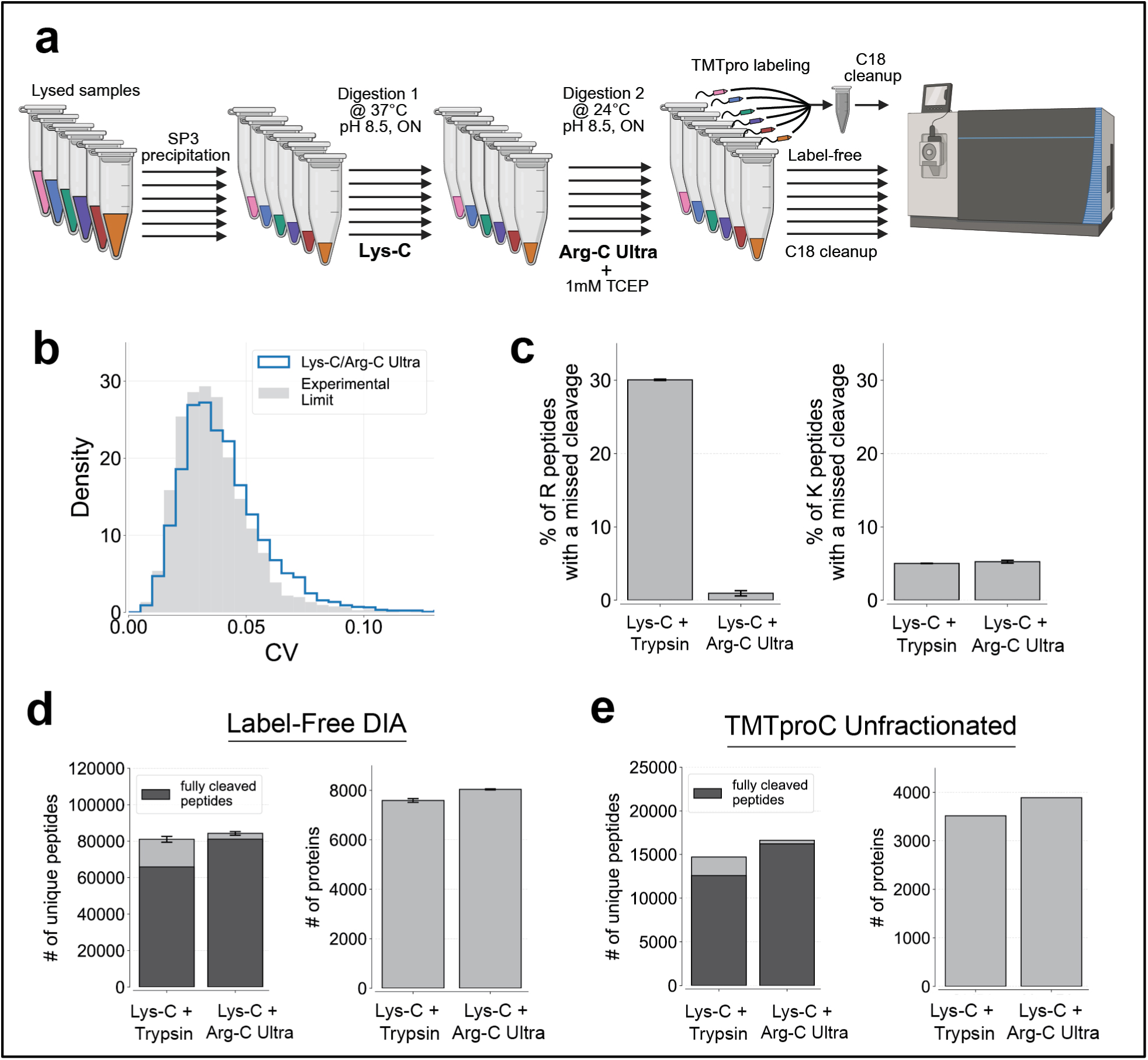
Lys-C and Arg-C Ultra synergistic digestion is highly reproducible and improves proteome coverage for multiplexed and label-free proteomics workflows. **(a)** Schematic of Lys-C/Arg-C Ultra digestion workflow compatible with multiplexed and label-free proteomics. Bead-bound proteins are initially digested with Lys-C (40 ng/uL) in 20 mM EPPS (pH 8.5) at 37 °C overnight. TCEP was then added to 1 mM, followed by Arg-C Ultra at a 1:200 enzyme-to-protein ratio, with a second overnight digestion performed at 24 °C. Samples were either analyzed label-free or labeled with a unique TMTpro tag. After C18 cleanup, samples were analyzed with TMTproC or label-free DIA. Created with BioRender.com. **(b)** The Lys-C/Arg-C Ultra workflow is highly reproducible. To evaluate reproducibility, a single precipitated HeLa lysate was divided into 6 identical aliquots and digested independently. This was done with both the Lys-C/Trypsin protocol and the Lys-C/Arg-C Ultra protocol. For comparison, we premixed the six TMTpro tags and labeled a single pool of HeLa peptides (from Lys-C/Arg-C Ultra digest) with this mixture (grey). Samples were analyzed by TMTproC (unfractionated). This distribution represents the experimental limit of reproducibility by eliminating variability due to sample preparation, thus revealing only the variability that arises from the data acquisition process, such as limited ion statistics. A stringent signal-to-noise cutoff of 852 (2+) and 2206 (3+) was used. **(c)** Lys-C/Arg-C Ultra digestion minimizes missed cleavages at arginine residues while maintaining high efficiency at lysine sites. When applied to DIA workflows on an Orbitrap Ascend with 200 ug loaded, missed cleavages were reduced from ∼30% to <1% for peptides where the first basic amino acid is arginine (R), while remaining <5% for peptides where the first basic amino acid is lysine (K), compared to Lys-C/Trypsin. Error bars represent ±1 standard deviation from n = 3 replicates (digested independently). **(d)** Lys-C/Arg-C Ultra digestion improves peptide and protein identifications for label-free DIA proteomics. From the same samples in (c), Lys-C/Arg-C Ultra digestion increased the number of unique, fully cleaved peptides by 23% and protein identifications by 6% compared to Lys-C/Trypsin digestion. **(e)** Lys-C/Arg-C Ultra digestion improves peptide and protein identifications for multiplexed proteomics. From the samples described in (b), Lys-C/Arg-C Ultra digestion increased the number of unique, fully cleaved peptides by 29% and protein identifications by 11% compared to Lys-C/Trypsin digestion. A sum signal-to-noise ratio cutoff of 136 (2+) and 353 (3+) was applied, corresponding to a 10% coefficient of variation (CV) from ion statistics^37,38^.

To assess the reproducibility and cleavage efficiency of this protocol, we divided precipitated HeLa lysate into six identical aliquots and performed digestion separately on each aliquot using our new Lys-C/Arg-C Ultra protocol. Each sample was labeled with a unique TMTpro tag and combined after labeling for analysis (Figure 5a). To establish an experimental limit for reproducibility, we also created a control experiment where TMTpro reagents were pre-mixed and added to a single pool of HeLa peptides, eliminating any variation due to sample preparation, including digestion. Comparison of the six separately digested samples with this experimental limit revealed that the digestion protocol achieved reproducibility approaching the experimental noise limit imposed by the experiment, particularly ion statistics (Figure 5b), demonstrating excellent technical precision. The impact on proteome coverage was substantial across multiple analytical platforms. Application of the Lys-C/Arg-C Ultra protocol to label-free data-independent acquisition (DIA) achieved >99% arginine cleavage efficiency and 95% lysine cleavage efficiency, representing a dramatic reduction in arginine missed cleavages from ∼30% to <1% compared to our standard Lys-C/Trypsin protocol (Figure 5c). This improved digestion efficiency translated directly to enhanced proteome coverage, yielded a 23% increase in unique, fully cleaved peptides and a 6% increase in protein identifications (Figure 5d). The benefits extended to TMTpro-based multiplexed proteomics, with the Lys-C/Arg-C Ultra protocol increasing the number of unique, fully cleaved peptides by 29% and protein identifications by 11% compared to standard Lys-C/Trypsin digestion when analyzed with TMTproC (Figure 5e). These results demonstrate the protocol’s broad applicability across diverse mass spectrometry acquisition strategies.

## Conclusion

Efficient and reproducible proteome digestion remains a central bottleneck in quantitative mass- spectrometry workflows. By systematically optimizing the biochemical conditions for Arg-C Ultra and Lys-C and then combining them in a sequential, redox-tuned protocol, we eliminated virtually all arginine missed cleavages (<1%) while preserving the high lysine-cleavage efficiency already afforded by Lys-C. Implementing this two-step strategy increased the number of unique, fully cleaved peptides by up to 29% and protein identifications by 6–11% across both label-free DIA and TMTpro multiplexed analyses. These benefits translated directly into high quantitative precision, approaching the limit imposed by ion statistics. Importantly, the workflow requires only modest adjustments to standard SP3-based sample preparation, uses affordable reagents (1 mM TCEP), and is compatible with existing acquisition methods and data-analysis pipelines. Together, these advances solve the persistent challenge of arginine missed cleavages in bottom-up proteomics, enabling deeper and more reliable proteome coverage.

**Figure S1.**
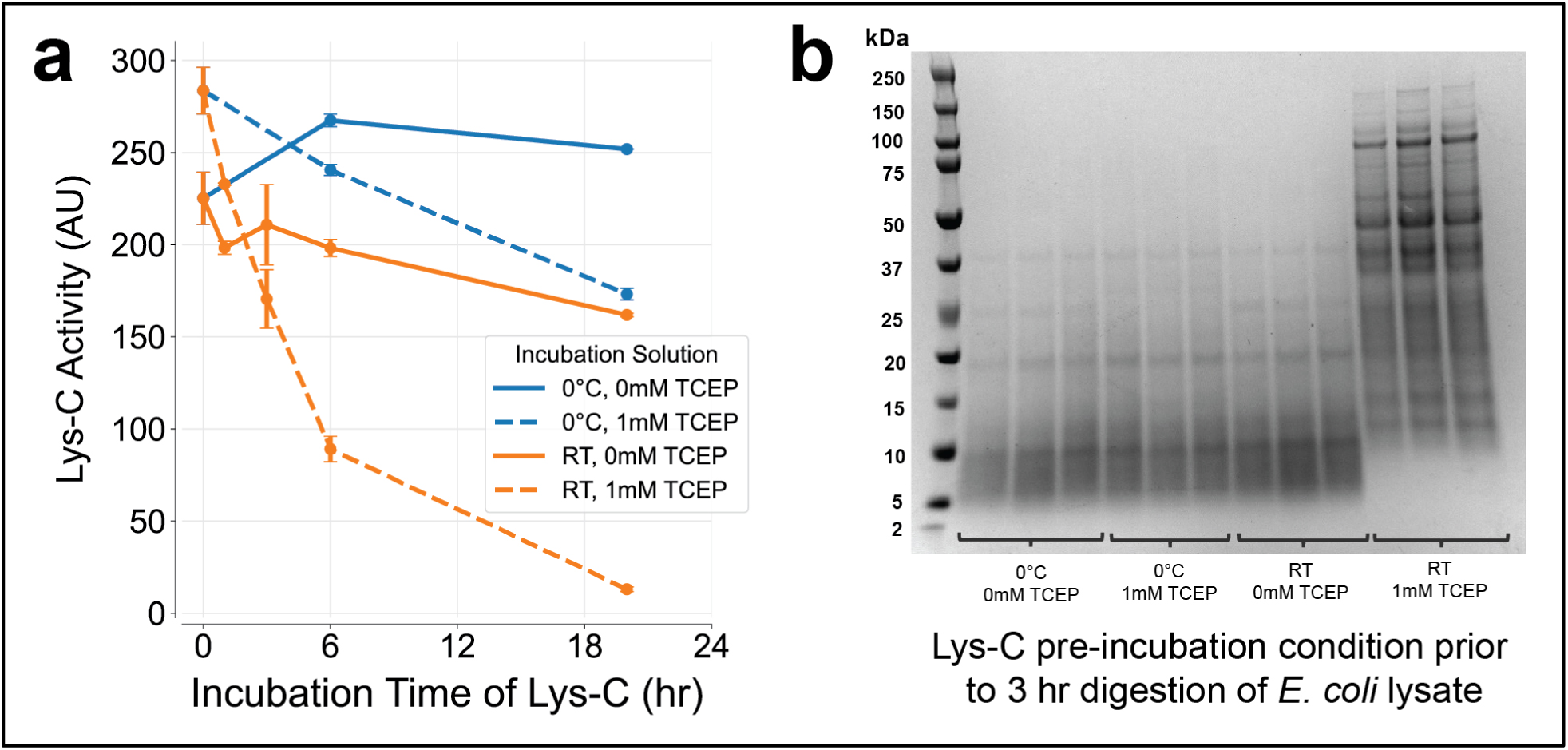
Validation that 1 mM TCEP Inhibits Lys-C Activity. **(a)** TCEP exposure strongly reduces Lys-C activity, measured via UV-Vis. Lys-C was incubated in 20 mM EPPS pH 8.5 with or without 1 mM TCEP at the indicated temperatures. Activity over time was measured using UV–Vis as in Figure 4. Exposure to TCEP at room temperature led to a >10-fold reduction in enzymatic activity after 20 hours. Error bars represent ±1 standard deviation from n=3 replicates. **(b)** Pre-incubation with 1mM TCEP renders Lys-C unable to digest lysate. Lys-C was pre-incubated in the indicated solutions overnight, then added to precipitated *E. coli* lysate for a 3-hour digestion at room temperature. The supernatant was collected, and proteins still bound to the beads were additionally released. Coomassie staining shows that Lys-C fails to digest the lysate when pre-incubated with 1 mM TCEP at room temperature.

**Figure S2.**
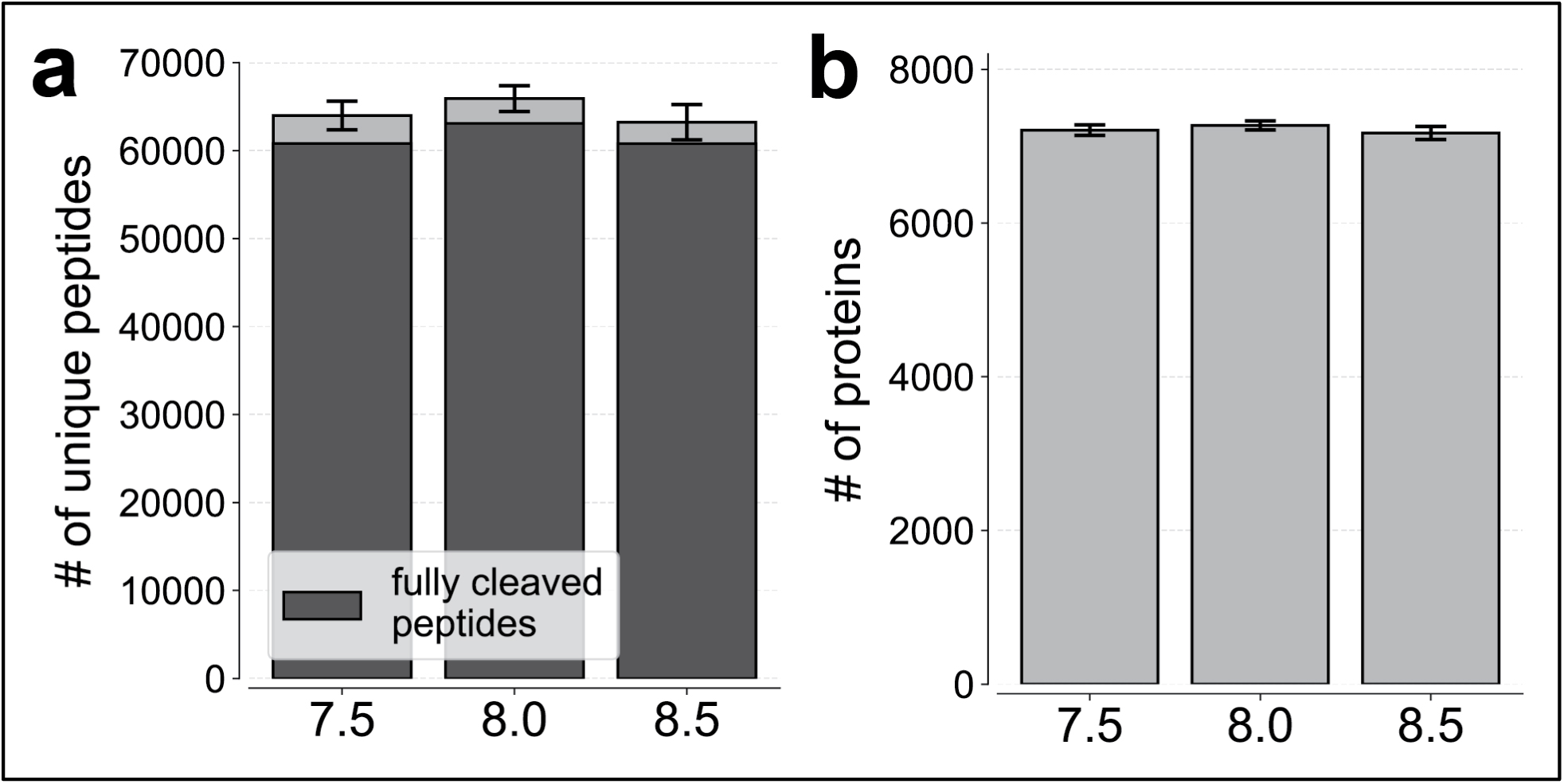
Effect of pH on digestion with the sequential Lys-C/Arg-C Ultra workflow. **(a)** Number of unique peptides. Bead-bound proteins were digested overnight with Lys-C (40 ng/uL) in 20 mM EPPS at pH 7.5, 8.0, or 8.5 (37 °C). TCEP (1 mM) and Arg-C Ultra (1:200 enzyme-to-protein) were then added for a second overnight digestion at 24 °C. All pH conditions performed similarly. Error bars represent ±1 standard deviation from n=3 replicates (digested independently). Data were acquired by label-free DIA on an Orbitrap Ascend. **(b)** Number of proteins. All pH conditions performed similarly

## Materials and Methods

### Cell Culture, Harvest, and Lysis

To generate HeLa cell pellets, HeLa S3 cells (ATCC, CCL-2.2) were cultured in Dulbecco’s Modified Eagle Medium (Corning, 10-013-CV) supplemented with 10% fetal bovine serum (Cytiva, SH30071.03) and 1× penicillin–streptomycin (Gibco, 15240-062) at 37 °C in a humidified incubator with 5% CO₂. Upon reaching ∼75% confluency, cells were split across twenty 100-mm cell culture-treated dishes. Once passaged cells reached ∼75% confluency again, cells were washed once with phosphate-buffered saline (Corning, 21-040-CM), detached using Trypsin-EDTA (Gibco, 25200-056), and pooled into 50 mL conical tubes. Suspensions were filtered through 70 um nylon strainers and adjusted to 1–2 × 10⁶ cells/mL based on cell counts from a trypan blue exclusion assay. 1 mL aliquots of the suspension were transferred into 1.5 mL microcentrifuge tubes, pelleted by centrifugation at 400×g for 5 minutes, washed twice with HEPES-buffered saline (50 mM HEPES, 140 mM NaCl, 1.5 mM Na₂HPO₄, pH 7.2), and centrifuged again to remove residual wash buffer by pipetting. The resulting pellets were flash-frozen in liquid nitrogen and stored at −80°C for downstream proteomics.

To generate *E.coli* cell pellets, K-12 MG1655 *E. coli* was plated from a glycerol stock. A single colony was inoculated overnight in MOPS minimal media. The media consisted of 40 mM MOPS buffer (Teknova, M2120) supplemented with 0.2% (w/v) glucose (Sigma, G8270), 9.5 mM ammonium chloride (NH₄Cl, Sigma, A9434), and 1.32 mM dipotassium phosphate (K₂HPO₄, Sigma, P3786), which were added separately. The next day, *E. coli* was diluted a minimum of 1:10 in MOPS minimal media, allowed to grow to no more than 0.4 OD₆₀₀, diluted again a minimum of 1:10, then grown to 0.4 OD₆₀₀, rapidly filtered with 0.2 um 90 mm cellulose acetate filter, scraped, and flash frozen.

For cell lysis, both HeLa and E. coli pellets were resuspended in lysis buffer containing 2% SDS, 50 mM HEPES (pH 7.2), and 1x cOmplete Mini, EDTA-free Protease Inhibitor Cocktail (Roche, 11836170001) to ∼1 ug/uL concentration. A needle sonicator was used with sonication for 30 seconds at 50% amplitude using direct sonication tips, followed by placement on ice for 30 seconds. This sonication/ice cycle was repeated 5 times. For HeLa lysate, following sonication, samples were heated to 95 °C for 20 minutes. Both lysates were then centrifuged at 20,000g for 15 minutes, and the supernatant was used for downstream processing.

### Proteomics Sample Preparation

Lysates were reduced with 5 mM DTT (Thermo Scientific, 165680050) for 20 minutes at 60°C, cooled to room temperature, alkylated with 20 mM N-ethylmaleimide (Thermo Scientific, 23030) for 20 minutes at room temperature, and quenched with 10 mM DTT for 10 minutes at room temperature. Proteins were precipitated using the SP3 protocol^16^ with a 1:1 mix of Sera-Mag SpeedBeads (Cytiva, 65152105050250 and 45152105050250). Beads were washed 3x with LC-MS grade water. A magnetic rack (DynaMag-2, Invitrogen™) was used to capture beads and/or bead-bound protein and remove supernatants. Beads were added to lysates at a 15:1 bead-to-protein mass ratio. Ethanol was added to a final concentration of 50% (v/v), and samples were incubated at room temperature at 1400 rpm for 1 hour on a thermomixer to initiate protein binding. Bead-bound proteins were washed 3x with 80% ethanol (v/v), with thermomixing and bath sonication as needed to ensure resuspension. Finally, bead-bound proteins were resuspended in the appropriate solution for digestion at ∼1 ug protein content/uL.

### Lys-C/Trypsin Digestions

Bead-bound protein was resuspended in 2 M GuHCl, 10 mM EPPS (pH 8.5) and Lys-C (Wako, Fisher Scientific, NC9242798) at 20 ng/uL to digest overnight (∼16 hr) at room temperature with thermomixing at 1400 rpm. Subsequently, the samples were diluted to 0.5 M GuHCl using 10 mM EPPS (pH 8.5) and digested with 20 ng/uL Lys-C and 10 ng/uL sequencing-grade Trypsin (Promega, V5113) at 37 °C overnight with thermomixing at 1400 rpm.

### Lys-C/Arg-C Ultra Digestions

Bead-bound protein was resuspended in 20 mM EPPS (pH 8.5) and Lys-C at 40 ng/uL to digest overnight at 37 °C with thermomixing at 1400 rpm. Subsequently, TCEP (VWR, K831-2G) was added to 1 mM followed by Arg-C Ultra at a 1:200 enzyme to protein content ratio by weight and digested overnight at room temperature with thermomixing at 1400 rpm.

After digestion, samples were placed on a magnetic rack, the supernatant was collected, and the beads were resuspended by pipette-mixing in the respective digestion buffer (0.5 M GuHCl, 10 mM EPPS or 20 mM EPPS) and placed back on the magnetic rack. This supernatant was combined with the previous supernatant and vacuum evaporated. For TMTpro labeling, peptides were resuspended in 200 mM EPPS pH 8.0 at ∼1 ug/uL concentration. For label-free, peptides were resuspended in 5% phosphoric acid. Samples were centrifuged at 20,000g for 15 minutes to remove residual beads.

### TMTpro labeling and desalting

For TMTpro labeling, peptides were incubated with TMTpro reagents (Thermo Fisher, A52045) at a 7.5:1 tag-to-peptide mass ratio for 2 hours at room temperature. The reaction was quenched with hydroxylamine (Sigma Aldrich, 467804) to a final concentration of 0.5% and incubated for an additional 30 minutes at room temperature. Labeled samples were pooled, vacuum evaporated to remove acetonitrile, and resuspended in 5% phosphoric acid. Peptides from both workflows were desalted using homemade StageTips packed with C18 material (Empore, 66883-U) as previously described^14^, and resuspended in 1% formic acid to a final concentration of ∼0.5 ug/uL prior to LC–MS analysis. Approximately 1 ug of peptides was analyzed unless otherwise stated.

### UHPLC Chromatography

Samples analyzed on the Orbitrap Fusion Lumos were run using an EASY-nLC 1200 system, while those analyzed on the Orbitrap Ascend used a Vanquish Neo UHPLC system (Thermo Fisher Scientific). The instruments were operated with Tune version 3.4 (Lumos) or 4.1 (Ascend). Peptides were separated on an IonOpticks Aurora Series emitter column (25 cm × 75 um ID, 1.6 um C18) maintained at 60 °C using a custom-built column oven. A nonlinear acetonitrile gradient was applied at a constant flow rate of 350 nL/min.

For the Orbitrap Lumos, solvent A consisted of 2% DMSO (LC–MS grade; Life Technologies), 0.125% formic acid (98%+; TCI America) in water (LC–MS grade; OmniSolv, VWR), and solvent B contained 80% acetonitrile (OmniSolv, MilliporeSigma), 2% DMSO, and 0.125% formic acid in water. For the Orbitrap Ascend, solvent A was 0.5% DMSO and 0.1% formic acid in water, and solvent B was 0.5% DMSO and 0.1% formic acid in acetonitrile.

For TMTpro samples, peptides were separated using a 90-minute gradient: 0–10% solvent B over 5 min, 10–26.4% B over 70 min, 26.4–100% B over 10 min, and held at 100% B for 5 min. For label-free DIA samples, a 70-minute gradient was used: 0–4.8% B over 5 min, 4.8–26.4% B over 50 min, 26.4–100% B over 10 min, and held at 100% B for 5 min. For label-free DDA samples, a 90-minute gradient was used. On the Orbitrap Ascend, the gradient was 0–4.8% B over 5 min, 4.8–26.4% B over 70 min, 26.4–100% B over 10 min, and held at 100% B for 5 min. On the Orbitrap Lumos, which used 80% acetonitrile in solvent B, the equivalent gradient was 0–6% B over 5 min, 6–33% B over 70 min, 33–100% B over 10 min, and held at 100% B for 5 min.

### TMTproC acquisition

TMTproC acquisitions were performed on the Orbitrap Ascend in positive ion mode using data-dependent MS2 analysis. RF lens level was set to 60%, and full MS1 scans were acquired in the Orbitrap at 120,000 resolution (at m/z 200), with an AGC target of 4.0e5, maximum ion injection time of 50 ms, and a scan range of m/z 350–1400. Wide quadrupole isolation was enabled. The maximum cycle time between full scans was set to 3 seconds. Monoisotopic precursor selection (“Peptide Mode”) was enabled, and isolated masses were dynamically excluded for 60 seconds within a ±10 ppm mass window. Isotope peaks and alternate charge states were excluded. MS2 scans were triggered only for z = 2 ions within m/z 500–1074 and z = 3 ions within m/z 350–1381, ensuring that complementary ion clusters would fall within the normal Orbitrap scan range. Additional intensity thresholds were applied: a minimum intensity of 3.0e5 for z = 2 precursors and 1.5e6 for z = 3. For MS2 acquisition, the AGC target was set to 7.5e4, with a maximum injection time of 59 ms. The quadrupole was used for precursor isolation with a 0.4 Th window, and CID fragmentation was applied at 30% collision energy (10 ms activation time, activation Q of 0.25). MS2 spectra were acquired in the Orbitrap at 30,000 resolution in normal scan range mode. All other instrument settings remained at default.

### Label-free DDA Acquisition

Label-free DDA acquisitions were primarily performed on the Orbitrap Ascend, except for the dataset shown in Figure 2e, which was acquired on the Orbitrap Lumos. On the Orbitrap Ascend, MS1 scans were acquired in positive ion mode using the Orbitrap at 120,000 resolution over a scan range of 350–1400 m/z with quadrupole isolation. The AGC target was set to 4.0e5, maximum injection time was 50 ms, and data were collected in centroid mode. Data-dependent MS2 scans were triggered on a 3-second cycle time. Monoisotopic precursor selection (“Peptide Mode”) was enabled, and isolated masses were dynamically excluded for 30 seconds within a ±10 ppm mass window. Isotope peaks and alternate charge states were excluded. Precursors were isolated using a 0.5 m/z window and HCD fragmentation was applied at a normalized collision energy of 35%. MS2 scans were acquired in the ion trap using Turbo scan mode over a range of 200–1400 m/z, with a normalized AGC target of 300% and a maximum injection time of 13 ms. Precursors with charge states 2–6 were selected for fragmentation. Label-free DDA acquisitions shown in Figure 2e were performed on an Orbitrap Fusion Lumos in positive ion mode. MS1 scans were acquired in the Orbitrap at 120,000 resolution over a scan range of 350–1750 m/z, with quadrupole isolation, an AGC target of 4.0e5, and a maximum injection time of 10 ms. Monoisotopic precursor selection (“Peptide Mode”) was enabled, and isolated masses were dynamically excluded for 60 seconds within a ±10 ppm mass window. Isotope peaks and alternate charge states were excluded. Precursor selection followed a charge- and mass-dependent decision tree: z = 2 precursors within 500–1750 m/z and z = 3–6 precursors within 600–1500 m/z were fragmented by CID at 30% collision energy, while z = 3–6 precursors within 350–600 m/z were fragmented by HCD at 24% collision energy. Precursors were isolated using a 0.5 m/z quadrupole window. MS2 spectra were acquired in the ion trap using Turbo scan mode over a range of 200–1400 m/z, with an AGC target of 1.0e4 and a maximum injection time of 35 ms. All data were acquired in centroid mode, and the cycle time was set to 1.5 seconds.

### TMTpro-SPS-MS3 Acquisition

TMTpro-SPS-MS3 acquisitions were performed on the Orbitrap Ascend using data-dependent MS3 analysis in positive ion mode. MS1 scans were acquired in the Orbitrap at 120,000 resolution (at m/z 200) over a scan range of 350–1500 m/z, with quadrupole isolation, an AGC target of 4.0e5, and a maximum injection time of 50 ms. RF lens voltage was set to 60%, and data were collected in centroid mode. Monoisotopic precursor selection was enabled, and dynamic exclusion was applied after a single occurrence for 60 seconds within a ±10 ppm mass tolerance. Isotope peaks and alternate charge states were excluded. Precursors with charge states 2–6 were selected for fragmentation and isolated using a 0.5 m/z window. MS2 scans were performed in the ion trap using CID at 35% normalized collision energy, with an AGC target of 1.0e4, maximum injection time of 50 ms, and Turbo scan mode. For MS3 acquisition, synchronous precursor selection (SPS) was enabled, selecting 5 fragment ions per precursor. SPS-MS3 scans were acquired in the Orbitrap at 45,000 resolution using HCD fragmentation at 45% collision energy. The AGC target was set to 2.0e5 with a maximum injection time of 91 ms. An MS2 isolation window of 1.2 m/z and an MS3 isolation window of 2.0 m/z were used. MS3 scans were collected in centroid mode with a scan range of 110–400 m/z.

### Data Independent Acquisition

All DIA-MS analysis was conducted using an Orbitrap Ascend. Peptides were separated over a 70-minute LC gradient in positive ion mode. MS1 scans were acquired in the Orbitrap at a resolution of 120,000 with a scan range of 350–1350 m/z. A quadrupole was used for precursor isolation, with an AGC target of 4.00e5. Data were collected in profile mode. MS2 spectra were acquired using data-independent acquisition (DIA) with a precursor m/z range of 350–950. The method used 75 consecutive 8 m/z-wide isolation windows with 0 m/z overlap. Fragmentation was performed using higher-energy collisional dissociation (HCD) with a normalized collision energy of 30%. MS2 scans were recorded in the Orbitrap at 15,000 resolution, with an AGC target of 5.00e5 and a maximum injection time of 27 ms. Data were collected in centroid mode.

### Data Analysis

Label-free DDA and TMTpro multiplexed data were analyzed using the Gygi Lab GFY (Core Version 3.12) software licensed from Harvard. Thermo Fisher Scientific raw-files were converted to mzXML using ReAdW.exe (http://svn.code.sf.net/p/sashimi/code/). MS2 spectra were assigned using the SEQUEST algorithm, searching against the UniProt reference proteomes for Homo sapiens (SwissProt + TrEMBL, downloaded August 7, 2016) supplemented with common contaminants (e.g., keratins, trypsin). For *E. coli* samples, the reference proteome UP000000625 from UniProt was used.

The target-decoy strategy was used to construct a second database of reversed sequences that were used to estimate the false discovery rate on the peptide level^39^. SEQUEST searches were performed using a 20-ppm precursor ion tolerance with the requirement that both N- and C-terminal peptide ends are consistent with the protease specificities of Lys-C and Trypsin. One missed cleavage was allowed. For high-resolution MS2 data (TMTproC) the fragment ion tolerance of the MS2 spectrum was set to 0.02 Da, whereas this value was set to 1 Da for low-resolution MS2 spectra acquired in the ion trap. TMTpro (+ 304.2071 Da) was set as a static modification on N-termini and lysine residues for labeled samples, and N-ethylmaleimide (+125.047679 Da) was set as a static modification on cysteine residues. Oxidation of methionine (+15.99492 Da) was set as a variable modification. The maximum number of variable modifications was set to 3. A peptide level MS2 spectral assignment false discovery rate of 1% was obtained by applying the target decoy strategy with linear discriminant analysis as described previously^40^. Peptides were assigned to proteins and a second filtering step to obtain a 1% FDR on the protein level was applied. Peptides that matched multiple proteins were assigned to the proteins with the most unique peptides^41^. Identification of complementary ion peaks, modelling of the isolation window, and deconvolution of the complementary peaks were performed as previously described^10^.

For DIA data, raw files were analyzed using DIA-NN version 2.2.0 with a library-free workflow. The precursor m/z range was set to 350–950. Oxidation of methionine was specified as the only variable modification (maximum of one variable modification per peptide). The default carbamidomethylation of cysteines was turned off, and N-ethylmaleimide alkylation was specified instead in the Additional Options pane. All samples were analyzed using the “Unrelated Runs” setting, meaning match-between-runs was disabled, as the study involved evaluating different experimental conditions rather than performing direct quantitative comparisons. All other parameters were left at default, including the allowance of one missed cleavage and default mass accuracy and scan window settings. A precursor-level false discovery rate (FDR) of 1.0% was applied by default. For protein-level filtering, protein groups with a PG.Q.Value ≤ 1% were retained. Spectral libraries were generated directly from the provided FASTA files.

All counts of unique peptides refer to distinct, stripped amino acid sequences (collapsed across charge states and modifications).

### UV-Vis spectrophotometry

Enzyme kinetics were measured using a Cary 300 UV-Vis spectrophotometer, equipped with Cary Dual Cell Peltier Accessory for temperature control. Stock reaction solutions were prepared for 2 mM Lysine (Nα-Benzyloxycarbonyl-L-lysine Thiobenzyl Ester, Calbiochem , 200274-100MG) or 1 mM Arginine (Nα-Benzoyl-L-arginine 4-nitroanilide, Sigma-Aldrich, B3133- 25MG) -like substrate in 20mM EPPS pH 8.5 (unless otherwise specified). Stocks were then diluted 10-fold with 20mM EPPS pH 8.5 to 0.2 mM for Lysine or 0.1 mM for Arginine-like substrate respectively. 2.5 mL and 400 uL of the solution was added to a 3 mL and 800 uL cuvette (1cm path length) respectively and blanked (zeroed). Samples were zeroed by placing the same solution mixture–minus the enzyme–in the rear of the UV-Vis. For Arg-C Ultra assays both cells were heated to 37° C and allowed to equilibrate for 2-3 minutes before the enzyme was added. For Lys-C assays measurements were taken at room temperature. Enzyme was then added to the 400 uL of reaction solution in the sample cell to a final concentration of 7 or 1.2 nM for Arg-C Ultra and Lys-C respectively. The absorption was then measured at 240 nm and 385 nm for Lysine and Arginine substrates respectively over 3 minutes. As several experimental assays were performed each one differed slightly as follows: For Arg-C Ultra experiments, 1mM of TCEP (VWR , K831-2G) was also added to the base solution mixture. For pH experiments the 20 mM EPPS used for diluting the 10x substrate stocks was pH’ed to a series of pH’s before diluting the concentrates. For temperature experiments, the samples were allowed to equilibrate for 4 minutes before enzyme was added and activity measured. For NHS-ester experiments, 20 mM EPPS at pH 8.0 was used to buffer reaction solutions. To make stock reaction solutions, various reducing agents were added to the final concentrations mentioned (0.1 mM TCEP, 0.4 mM TCEP, 0.5 mM TCEP, 0.7 mM TCEP, 1 mM TCEP, 1.3 mM TCEP, 5 mM TCEP, 1 mM BME, and 1 mM DTT). Solutions were pH’ed back to 8.0 after addition of reducing agents. These solutions were then blanked and zeroed as before and then TMTpro0 (ThermoFisher Scientific, A44518) was added to a final concentration of 96 nM and the absorption measured at 260 nm for 3 minutes.

### Visualization of protein structure

The structure of wild-type *Lysobacter enzymogenes* Lys-C endoproteinase (PDB: 4NSY) was visualized using PyMOL (Schrödinger, LLC). Only chain A was retained, and all non-protein heteroatoms were removed. Cysteines involved in disulfide bonds were shown as sticks and colored orange.

### Gel Electrophoresis and Staining

Protein samples were prepared under non-reducing conditions by resuspension in 1X Bolt LDS Sample Buffer (Thermo Fisher, B0007). Samples were vacuum evaporated to dryness prior to resuspension. Samples were then heated at 70 °C for 10 minutes. Proteins were separated by SDS-PAGE using NuPAGE 4–12% Bis-Tris Mini Gels (Invitrogen, NP0322BOX) in MES SDS Running Buffer (Invitrogen, NP0002), with electrophoresis performed at a constant voltage of 200 V.

Following electrophoresis, gels were removed from the cassette and fixed in a solution of 40% methanol, 10% acetic acid, and 50% deionized water for 10 minutes with gentle rocking. Gels were subsequently stained with Imperial Protein Stain (Thermo Fisher, 24615) for 1 hour with rocking, then destained in a solution of 10% methanol, 10% acetic acid, and 80% deionized water until the background was sufficiently clear.

To visualize protein digestion patterns shown in Figure 2f , 4e, and S1b, after the digestion supernatants were collected, the beads were boiled in 2% SDS at 95 °C for 30 min to release any remaining undigested proteins from beads. After boiling, samples were placed immediately on a magnetic rack, and the supernatant was collected. This was combined with the digestion supernatant and vacuum evaporated to dryness, then resuspended in sample buffer for gel-electrophoresis.

## Acknowledgement

This work was supported by NIH grant R35GM128813 (MW), NIH training grant T32GM007388, Ludwig Cancer Research (MW), Princeton’s Summer Undergraduate Research Program in Molecular and Quantitative & Computational Biology (VP, GB), and Summer Undergraduate Research Fellows in Chemistry program (CN). We also gratefully acknowledge support from the Princeton Catalysis Initiative (MW). The content of this article is solely the responsibility of the authors and does not necessarily represent the official views of the National Institutes of Health.

## References

1. Nesvizhskii, A. I. & Aebersold, R. Interpretation of Shotgun Proteomic Data. Molecular & Cellular Proteomics 4, 1419–1440 (2005).

2. Yates, J. R. The Revolution and Evolution of Shotgun Proteomics for Large-Scale Proteome Analysis. J Am Chem Soc 135, 1629 (2013).

3. Loziuk, P. L. et al. Understanding the role of proteolytic digestion on discovery and targeted proteomic measurements using liquid chromatography tandem mass spectrometry and design of experiments. J Proteome Res 12, 5820–5829 (2013).

4. Brownridge, P. & Beynon, R. J. The importance of the digest: Proteolysis and absolute quantification in proteomics. Methods 54, 351–360 (2011).

5. Leon, I. R., Schwammle, V., Jensen, O. N. & Sprenger, R. R. Quantitative assessment of in-solution digestion efficiency identifies optimal protocols for unbiased protein analysis. Molecular and Cellular Proteomics 12, 2992–3005 (2013).

6. Walmsley, S. J. et al. Comprehensive analysis of protein digestion using six trypsins reveals the origin of trypsin as a significant source of variability in proteomics. J Proteome Res 12, 5666 (2013).

7. Olsen, J. V., Ong, S. E. & Mann, M. Trypsin Cleaves Exclusively C-terminal to Arginine and Lysine Residues. Molecular & Cellular Proteomics 3, 608–614 (2004).

8. Sun, W., Wu, S., Wang, X., Zheng, D. & Gao, Y. A Systematical Analysis of Tryptic Peptide Identification with Reverse Phase Liquid Chromatography and Electrospray Ion Trap Mass Spectrometry. Genomics Proteomics Bioinformatics 2, 174–183 (2004).

9. Guthals, A. & Bandeira, N. Peptide Identification by Tandem Mass Spectrometry with Alternate Fragmentation Modes. Molecular & Cellular Proteomics 11, 550–557 (2012).

10. Sonnett, M., Yeung, E. & Wühr, M. Accurate, Sensitive, and Precise Multiplexed Proteomics Using the Complement Reporter Ion Cluster. Anal Chem 90, 5032–5039 (2018).

11. Šlechtová, T., Gilar, M., Kalíková, K. & Tesařová, E. Insight into Trypsin Miscleavage: Comparison of Kinetic Constants of Problematic Peptide Sequences. Anal Chem 87, 7636–7643 (2015).

12. Gershon, P. D. Cleaved and missed sites for trypsin, Lys-C, and Lys-N can be predicted with high confidence on the basis of sequence context. J Proteome Res 13, 702–709 (2014).

13. Glatter, T. et al. Large-scale quantitative assessment of different in-solution protein digestion protocols reveals superior cleavage efficiency of tandem Lys-C/trypsin proteolysis over trypsin digestion. J Proteome Res 11, 5145–5156 (2012).

14. Gupta, M., Sonnett, M., Ryazanova, L., Presler, M. & Wühr, M. Quantitative Proteomics of Xenopus Embryos I, Sample Preparation. Methods Mol Biol 1865, 175 (2018).

15. Li, Q., Feng, Y., Tan, M. J. & Zhai, L. H. Evaluation of Endoproteinase Lys-C/Trypsin Sequential Digestion Used in Proteomics Sample Preparation. Chinese Journal of Analytical Chemistry 45, 316–321 (2017).

16. Hughes, C. S. et al. Single-pot, solid-phase-enhanced sample preparation for proteomics experiments. Nat Protoc 14, 68–85 (2019).

17. Johnson, A. N. T. et al. Sensitive and Accurate Proteome Profiling of Embryogenesis Using Real-Time Search and TMTproC Quantification. Molecular & Cellular Proteomics 24, 100899 (2025).

18. Poulsen, J. W., Madsen, C. T., Young, C., Poulsen, F. M. & Nielsen, M. L. Using guanidine-hydrochloride for fast and efficient protein digestion and single-step affinity-purification mass spectrometry. J Proteome Res 12, 1020–1030 (2013).

19. Schmitz, E. M. H. et al. Optimizing charge state distribution is a prerequisite for accurate protein biomarker quantification with LC-MS/MS, as illustrated by hepcidin measurement. Clin Chem Lab Med 56, 1490–1497 (2018).

20. Jeong, K., Kim, S. & Bandeira, N. False discovery rates in spectral identification. BMC Bioinformatics 13 Suppl 16, 1–15 (2012).

21. Meyer, J. G. & Komives, E. A. Charge state coalescence during electrospray ionization improves peptide identification by tandem mass spectrometry. J Am Soc Mass Spectrom 23, 1390–1399 (2012).

22. Chen, Z., Potempa, J., Polanowski, A., Wikstrom, M. & Travis, J. Purification and characterization of a 50-kDa cysteine proteinase (gingipain) from Porphyromonas gingivalis. Journal of Biological Chemistry 267, 18896–18901 (1992).

23. Pike, R., McGraw, W., Potempa, J. & Travis, J. Lysine- and arginine-specific proteinases from Porphyromonas gingivalis. Isolation, characterization, and evidence for the existence of complexes with hemagglutinins. Journal of Biological Chemistry 269, 406– 411 (1994).

24. Nakayama, K., Kadowaki, T., Okamoto, K. & Yamamoto, K. Construction and characterization of Arginine-specific cysteine proteinase (Arg-gingipain)-deficient mutants of Porphyromonas gingivalis: Evidence for significant contribution of Arg-gingipain to virulence. Journal of Biological Chemistry 270, 23619–23626 (1995).

25. Potempa, J. et al. Comparative properties of two cysteine proteinases (gingipains R), the products of two related but individual genes of Porphyromonas gingivalis. Journal of Biological Chemistry 273, 21648–21657 (1998).

26. Eichinger, A. et al. Crystal structure of gingipain R: an Arg-specific bacterial cysteine proteinase with a caspase-like fold. EMBO J 18, 5453 (1999).

27. Labrou, N. E. & Rigden, D. J. The structure-function relationship in the clostripain family of peptidases. Eur J Biochem 271, 983–992 (2004).

28. Longarini, E. J. & Matić, I. Preserving ester-linked modifications reveals glutamate and aspartate mono-ADP-ribosylation by PARP1 and its reversal by PARG. Nat Commun 15, 4239 (2024).

29. Greene, R. F. & Pace, C. N. Urea and Guanidine Hydrochloride Denaturation of Ribonuclease, Lysozyme, α-Chymotrypsin, and b-Lactoglobulin. Journal of Biological Chemistry 249, 5388–5393 (1974).

30. Herbert, B. Advances in protein solubilisation for two-dimensional electrophoresis. Electrophoresis 20, 660–663 (1999).

31. Proc, J. L. et al. A quantitative study of the effects of chaotropic agents, surfactants, and solvents on the digestion efficiency of human plasma proteins by trypsin. J Proteome Res 9, 5422–5437 (2010).

32. Ren, D. et al. An improved trypsin digestion method minimizes digestion-induced modifications on proteins. Anal Biochem 392, 12–21 (2009).

33. Alexandrova, A. N. & Jorgensen, W. L. Why Urea Eliminates Ammonia Rather Than Hydrolyzes in Aqueous Solution. J Phys Chem B 111, 720 (2007).

34. Kollipara, L. & Zahedi, R. P. Protein carbamylation: In vivo modification or in vitro artefact? Proteomics 13, 941–944 (2013).

35. Hoeven, L. van der, Lechner, M., Hernandez-Rollan, C., Batth, T. S. & Olsen, J. V. Comparative Analysis of Lysine-Specific Peptidases for Optimizing Proteomics Workflows. bioRxiv 2024.10.18.619105 (2024) doi:10.1101/2024.10.18.619105.

36. Miron, T. & Wilchek, M. A spectrophotometric assay for soluble and immobilized N-hydroxysuccinimide esters. Anal Biochem 126, 433–435 (1982).

37. Peshkin, L., Gupta, M., Ryazanova, L. & Wühr, M. Bayesian Confidence Intervals for Multiplexed Proteomics Integrate Ion-statistics with Peptide Quantification Concordance,. Molecular & Cellular Proteomics 18, 2108–2120 (2019).

38. Johnson, A., Stadlmeier, M. & Wühr, M. TMTpro Complementary Ion Quantification Increases Plexing and Sensitivity for Accurate Multiplexed Proteomics at the MS2 Level. J Proteome Res 20, 3043–3052 (2021).

39. Elias, J. E. & Gygi, S. P. Target-decoy search strategy for increased confidence in large-scale protein identifications by mass spectrometry. Nat Methods 4, 207–214 (2007).

40. Huttlin, E. L. et al. A tissue-specific atlas of mouse protein phosphorylation and expression. Cell 143, 1174–1189 (2010).

41. Savitski, M. M., Wilhelm, M., Hahne, H., Kuster, B. & Bantscheff, M. A scalable approach for protein false discovery rate estimation in large proteomic data sets. Molecular and Cellular Proteomics 14, 2394–2404 (2015).

